# Dietary Exposure to Purified Gliadins and Avenins: Effects on Intestinal and Immune Responses in BALB/c Mice

**DOI:** 10.64898/2026.10.08.757589

**Authors:** Michaela Šťastná, Adam Norek, Peter Makovický, Edita Jeklová, Petra Straková, Matěj Hrunka, Petr Jabandžiev, Lubomír Janda

## Abstract

Dietary cereal prolamins differ in composition and gastrointestinal processing, but their shared and protein-source-specific effects on intestinal homeostasis remain incompletely defined. BALB/c mice were fed for five weeks with diets in which 70% of total protein was supplied by vital wheat gluten, purified gliadin, or purified avenin, alongside casein-fed controls. Intestinal morphology, antigen-specific plasma immunoglobulin G and A (IgG, IgA), mucosal gene expression, and small-intestinal proteomics were analysed. No overt histopathological injury or generalised inflammatory response was detected in any group. Gliadin and avenin both increased ileal villus length; avenin additionally elicited the broadest and most frequent prolamin-specific IgG response, associated with reduced intestinal interleukin-10 expression in responding animals. Proteomic profiling revealed a shared structural-metabolic signature alongside protein-specific changes in lipid handling, junctional regulation, and epithelial immune signalling. These findings indicate that wheat-and oat-derived prolamins differentially influence intestinal homeostasis and antigen recognition even at matched dietary doses, informing their evaluation as functional food ingredients.

**Graphical abstract:** 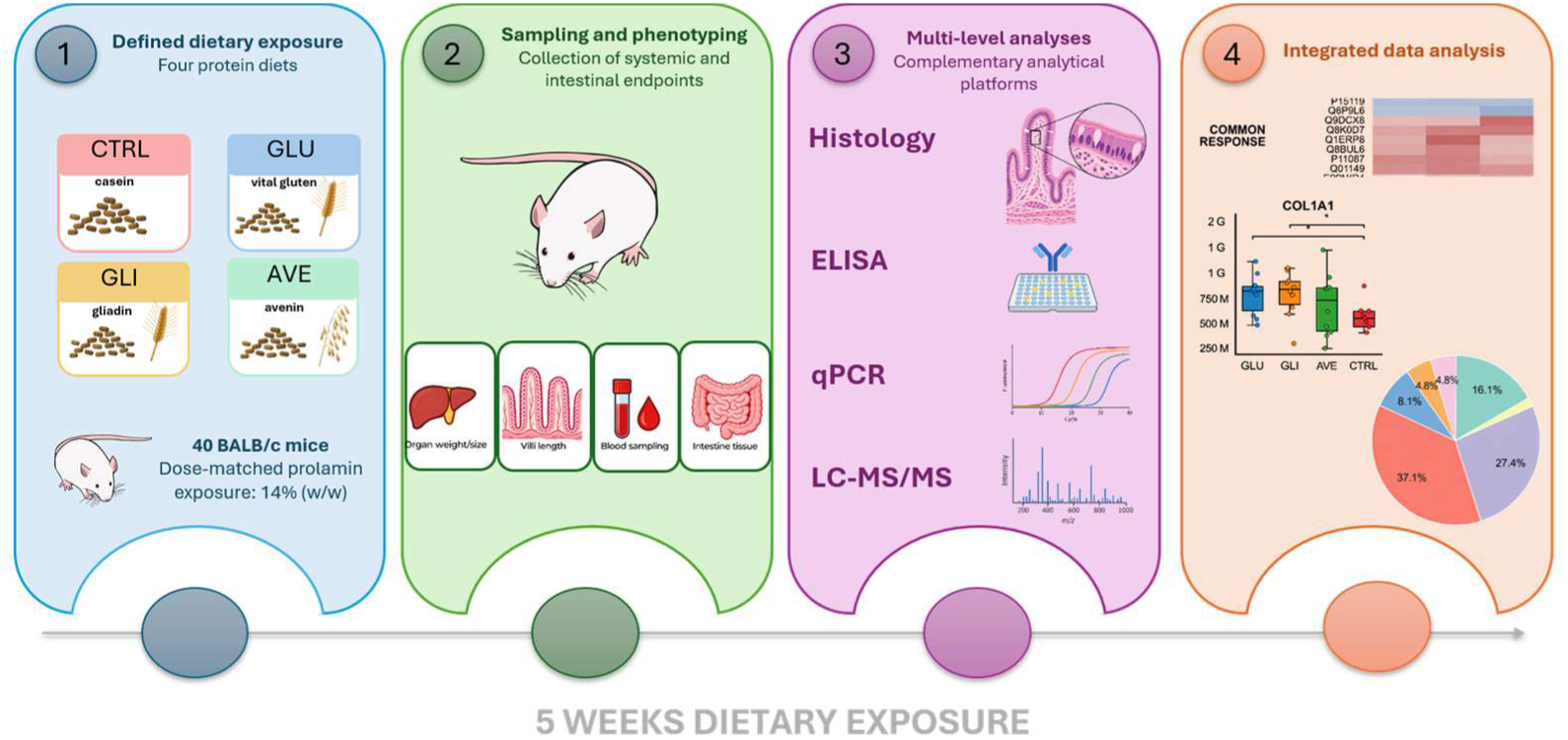

## 1. Introduction

Cereal prolamins represent a major class of dietary storage proteins with distinct biochemical properties and the capacity to interact with the intestinal mucosal environment (Holding, 2014; P. R. Shewry & Halford, 2002). Among these, wheat-derived gliadins are well-established drivers of immune activation, frequently inducing adaptive responses and characteristic mucosal shifts in sensitive individuals (Cenni et al., 2023; Fasano & Catassi, 2012; Sapone et al., 2012). In contrast, oat-derived prolamins (avenins) are generally considered less immunogenic and uncontaminated gluten-free oats are well tolerated by most individuals consuming gluten-free diets (Guttormsen et al., 2008; Hoffmanová et al., 2019; Janatuinen et al., 1995; Kaukinen et al., 2013). Nevertheless, evidence of avenin-specific immune responses in selected individuals indicates that avenins should not necessarily be regarded as biologically inert at the intestinal interface (Lundin et al., 2003; Tuire et al., 2012). Whether gliadins and avenins differ in their capacity to modulate intestinal homeostasis independently of overt tissue injury remains insufficiently understood.

Prolamins are traditionally classified by the Osborne system based on their solubility in aqueous alcohols. Wheat gluten comprises monomeric gliadins (α/β, γ, and ω subtypes) and polymeric glutenins and is characterised by a high abundance of proline-and glutamine-rich sequences (P. Shewry, 2023). These structural features contribute to incomplete gastrointestinal proteolysis and the persistence of relatively protease-resistant peptides. Avenins constitute a substantially smaller proportion of oat grain proteins and differ from wheat gliadins in amino-acid composition, solubility, and susceptibility to gastrointestinal proteolysis (Gilissen et al., 2016). Such biochemical differences may influence not only peptide persistence but also antigen availability and interactions with the intestinal epithelium and mucosal immune system.

This distinction is particularly relevant because comparisons between wheat and oat prolamins under physiological dietary conditions are inherently influenced by different antigenic loads. Wheat typically contains substantially higher levels of prolamins than oats, often by a factor of five or more (Kosová et al., 2020). Consequently, it is difficult to distinguish intrinsic protein-specific properties from effects related simply to the amount of antigen reaching the intestine. A dose-matched comparison of purified gliadins and avenins therefore provides an experimental approach to separate qualitative differences between these cereal proteins from quantitative exposure-dependent effects.

The intestinal epithelium is highly responsive to changes in luminal composition (Le Gall et al., 2019). Food proteins and their digestion products may influence epithelial renewal, proteostasis, antigen processing and transport, and mucosal immune regulation without necessarily inducing classical inflammation or structural damage. Maintenance of tolerance to dietary antigens depends on coordinated epithelial and immune mechanisms, including tolerogenic antigen-presenting cells, regulatory T-cell responses, and immunoregulatory cytokines such as IL-10 and TGF-β (Pabst & Mowat, 2012; Weiner, 2000). Consequently, antigen-specific systemic immune responses may provide evidence of altered host recognition even when intestinal morphology remains preserved. Examining intestinal proteomic networks together with local immune regulation and systemic antibody responses may therefore reveal subtle biological effects of dietary proteins that are not captured by conventional histopathological assessment alone.

In the present study, we used a five-week dietary intervention in BALB/c mice to compare the effects of purified wheat gliadins and oat avenins administered at equivalent high dietary concentrations, alongside gluten and control diets. By integrating intestinal morphometry, proteomic profiling, mucosal gene expression, and antigen-specific antibody responses, we investigated whether structurally distinct cereal prolamins differentially modulate intestinal homeostasis and systemic immune recognition in the absence of overt inflammatory pathology. We hypothesised that dose-matched exposure to gliadins and avenins would reveal protein-specific and host-dependent differences in epithelial homeostatic networks, antigen handling, and immune regulation that are obscured under conditions of unequal dietary exposure.

## 2. Materials and Methods

### Prolamin preparation

Vital gluten (Lantmännen Reppe AB, Lidköping, Sweden) was used for gliadin isolation. In one run, 4 kg of vital gluten was resuspended in 34 L of 80% (v/v) ethanol, stirred well and incubated overnight in a closed container. The next day, the suspension was centrifuged (5,000g, 20 min, RT), and the supernatant was carefully transferred to a new container. Gliadins were precipitated at 4 °C according to Tanner et al.,(2019). The suspension was then centrifuged (5,000 g, 40 min, 4 °C), and the resulting pellet was placed on a tray and dried at 60 °C for 2 days, then further dried at 80 °C. The dried gliadins were subsequently ground using a mill (IKA, Staufen, Germany). Avenins were isolated from oat flour of the Atego variety (Selgen as., Sibřina, Czech Republic), a cultivar for which the avenin-encoding genes have been sequenced (Leišová-Svobodová et al., 2022). In one run, 12 kg of oats was resuspended in 34 L of 60% ethanol, stirred well, and incubated overnight in a closed container. The following steps were the same as for gliadin isolation.

Isolated avenins and gliadins were analysed at the Food Research Institute (Prague, Czech Republic) for total protein (Kjeldahl method) and fat content (gravimetric analysis after acid hydrolysis and chloroform extraction). Samples were further characterized by mass spectrometry at Central European Institute of Technology (CEITEC MU, Brno, Czech Republic). Proteins were partially dissolved in 0.1% DDM (60 min, 95 °C), reduced with DTT (30 min, 56 °C), alkylated with IAA (30 min, 25 °C), digested with trypsin (4 h, 37 °C), and analyzed by LC-MS/MS (RSLCnano system, Thermo Fisher Scientific; Impact II, Bruker). MS/MS data were searched using MASCOT against the UniRef100 database.

Protein isolates were also analyzed by western blot using monoclonal antibodies G12 (Biomedal, Sevilla, Spain) and GLD7 (Sigma-Aldrich, Burlington, Massachusetts, USA). Vital wheat gluten, commercial gliadin (Sigma-Aldrich, Burlington, Massachusetts, USA), and the isolated proteins were dissolved in Laemmli buffer at 100 ng and 1 µg concentrations, incubated (15 min, 95 °C), and separated on 12.5% SDS-PAGE gels. Proteins were transferred to PVDF membranes and probed with G12 (dilution 1:40,000 of G12 antibody, 100 ng of each antigen) or GLD7 (dilution 1:15,000 of GLD7 antibody, 1 µg of each antigen). A goat anti-mouse IgG-Fc HRP-conjugated secondary antibody (Bethyl, Montgomery, Texas, USA) was used at 1:20,000 dilution and used at this concentration for G12 and GLD7 monoclonal antibody.

### Animals

All animal experiments complied with ARRIVE guidelines and were conducted in accordance with EU Directive 2010/63 and institutional regulations. The study was approved by Ministry of Agriculture, Department of Animal Commodities and Animal Welfare, approval no. MZe 2340. Forty BALB/c mice (20 males and 20 females), fourteen weeks old, were obtained from AnLab (Prague, Czech Republic). The animals were group-housed (2–5 animals of the same sex per cage) in a room with a 12-hour light cycle and a controlled temperature (26 ± 2 °C). All mice had free access to food and water. The animals were weighed regularly each week. Although both male and female mice were included, the study was not designed or powered to assess sex-specific differences, and the results were therefore analysed collectively.

Before the start of the experiment, the animals were divided into four groups (5 males and 5 females in each group) and fed four different experimental diets for 5 weeks. The first group was a control group fed a gluten-free diet (CTRL; AIN93G, E15712-04, growth and reproduction); the second group was a gluten group (GLU; Gluten-free AIN93G diet where 70% of total protein was replaced by vital gluten, 30% of protein content was casein); the third group was a gliadin group (GLI; Gluten-free AIN93G diet where 70% of total protein was replaced by gliadins, 30% of protein content was casein); and the last group was on an avenin diet (AVE; Gluten-free AIN93G diet where 70% of the total protein was replaced by avenins, 30% of protein content was casein). An amount corresponding to 70% of the total protein content represents 14% (w/w) of the diet. All diets were prepared by Ssniff (Soest, Germany) and all types of feed were nutritionally adequate and contained a balanced amount of nutrients recommended for rodents.

### Macroscopic and histological investigations

After five weeks, the mice were euthanized using inhalation anaesthesia (Isoflurane 1000 mg/g, Werfft, spol. s.r.o., Prague, Czech Republic) followed by cervical dislocation. The weight, length of the body and chest width were measured. A standard necropsy and macroscopic examination were performed. Internal organs, including the heart, liver, spleen, both kidneys, and, in male mice, both testes, were excised and weighed. In addition, the lengths of the small and large intestines were measured.

One sample from the duodenal, jejunal, and ileal sections, as well as one from the colon, was collected from each mouse. The material was fixed in 10% neutral buffered formalin solution (Sigma-Aldrich, Burlington, Massachusetts, USA) for 24 hours. Samples were routinely processed and embedded into paraffin blocks using an automated tissue processor (Leica, ASP6028, Leica Microsystems, Wetzlar, Germany). From three to five μm thick slices were cut from each sample using a rotary microtome (Leica RM2255, Leica Microsystems, Wetzlar, Germany) and tissue sections were placed on standard glass slides (Waldemar Knittel, Braunschweig, Germany). The slices were stained with hematoxylin-eosin (Bamed, Litvínovice, Czech Republic). The samples were described and evaluated in light-microscopic images using an optical microscope (Olympus BX46, Tokyo Japan). Objective assessment, achieved by the measurement of the height of villi and the depth of crypts, was carried out using NIS-Elements software, version 3.0 (Laboratory Imaging s.r.o., Prague, Czech Republic). In each sample, several areas were selected, in which a total of fifty villi and fifty crypts were measured.

### Assessment of specific IgG and IgA in plasma

Following euthanasia, blood samples were collected via cardiac puncture, and plasma was isolated by centrifugation. Antigen-specific IgG and IgA levels were quantified using an indirect ELISA. Microtiter plates were coated with 500 ng/well of each antigen (gluten, gliadin, avenin, and casein diluted in 0.05 M carbonate buffer, pH 9.6) and blocked with 3% bovine serum albumin (BSA). To assess potential cross-reactivity with dietary components, IgG responses against all antigens were evaluated across all experimental groups. Plasma samples were analysed in a serial dilution range (1:50 to 1:6,250). HRP-conjugated secondary antibodies were applied at dilutions of 1:5,000 (anti-IgG) and 1:2000 (anti-IgA). Absorbance was measured at 450 nm using a Synergy H1 microplate reader (BioTek, Santa Clara, CA, USA).

### RNA isolation and reverse transcription (RT)-qPCR

From intestinal homogenates, total RNA was extracted using phase separation procedure with bromanisole and purified using the NucleoSpin RNA Plus Mini Kit (Macherey-Nagel, Düren, Germany) according to the manufacturer’s instructions. RT of 1 μg of RNA was done using the Luna Script RT SuperMix Kit (NEB, Ipswich, Massachusetts, USA). qPCR was performed using Nanodrop II liquid dispenser (Innovadyne Technologies, Rohnert Park, California, USA) and LightCycler 480 instrument (Roche, Basel, Switzerland) according to the manufacturers’ instructions. One reaction contained: 10 pmol of each of paired gene specific primers (Generi Biotech, Hradec Kralove, Czech Republic; all sequences are in Supplementary Data C), 1.5 μL of QuantiTect SYBR Green PCR MasterMix (Qiagen, Darmstadt, Germany), and 4x diluted cDNA (0.5 μL). The primers were designed at the exon-exon junctions to avoid amplifying genomic DNA. Reference housekeeping gene (REF) TBP1 was determined using variability test among samples tested. Assuming primer efficiency ≥1.9, normalised gene expression based on quantification cycle (cq) values were calculated as 2-(CqGENE - CqREF) = 2(−ΔCt). Analysis of the melting temperature confirmed the specificity of amplicons using LightCycler 480 1.5.0.39 software (Roche Applied Science, Basel, Switzerland). Negative controls were set up in all parts of the gene expression assessment.

### Mass spectrometry

Proteins extracted from M. musculus intestine tissue were subjected to filter-aided sample preparation as described elsewhere (Wiśniewski et al., 2009). The resulting peptides were analysed by liquid chromatography-tandem mass spectrometry (LC-MS/MS) performed using UltiMate 3000 RSLCnano system (Thermo Fisher Scientific) online coupled with Orbitrap Exploris 480 spectrometer (Thermo Fisher Scientific). See the supplemental material section for full details regarding the analyses and data evaluation.

### Data processing and statistical analysis

Data processing and statistical analyses were performed in Python using pandas, NumPy, SciPy and Pingouin. All datasets were processed using a unified workflow to minimise the influence of individual sample variability and to keep the analysed animal set consistent across proteomic, ELISA, organ and intestinal morphometry data. Potentially biased samples were identified from the proteomic dataset by within-group rank-based quality control across retained proteins. Samples repeatedly occurring at extreme rank positions or showing an absolute rank-based Z-score > 3.0 were excluded, and the resulting exclusion mask was applied to downstream datasets.

For the broad descriptive proteomic overview, proteins identified by at least one peptide were assigned to composite GO-derived functional categories. For protein-level inferential analyses, the MS/MS dataset was filtered according to unique-peptide evidence, retaining proteins identified by at least two unique peptides. Median protein intensities were calculated for each experimental group. Proteins were selected for statistical testing when at least one experimental group differed from CTRL by at least 25%. Group differences were analysed using Welch’s one-way ANOVA, followed, when significant, by Games-Howell post-hoc comparisons of GLU, GLI and AVE against CTRL. Multiplicity-adjusted post-hoc p < 0.05 was considered statistically supported. IgG-positive and IgG-negative animals were compared using Welch’s two-sample t-test, with p < 0.05 considered significant.

ELISA, RT-qPCR, organ and intestinal morphometry data were analysed using the same general framework. Repeated measurements were averaged to one value per animal. For organ and intestinal morphometry variables, group-wise extreme values were removed using the 5th-95th percentile range before statistical testing. Targeted endpoints were analysed using Welch’s ANOVA followed, where appropriate, by Games-Howell comparisons against CTRL, with p < 0.05 considered significant.

## 3. Results and Discussion

### 3.1. Verification of the prolamin composition in dietary protein isolates

To ensure accurate interpretation of in vivo responses, the composition and immunological properties of the dietary protein isolates were characterised prior to administration. All preparations showed high protein content, reaching 74.9 g/100 g in vital wheat gluten, 95.2 g/100 g in purified gliadin, and 88.0 g/100 g in purified avenin, with minimal fat contamination (0.32–4.56 g/100 g), confirming their suitability as defined protein sources (Supplementary Data A).

Proteomic profiling via mass spectrometry (MS) confirmed that vital wheat gluten comprised both glutenins and prolamins, whereas the purified gliadin fraction was highly enriched in wheat prolamins with only minor co-isolated proteins. Similarly, the avenin preparation predominantly consisted of oat prolamins with negligible contamination, confirming a high degree of compositional specificity (Supplementary Data B).

Immunoblotting further characterised the antigenic properties of the isolates. The G12 antibody, which recognises immunodominant gliadin hexameric epitope QPQLPY (Morón et al., 2008), displayed high affinity for wheat-derived proteins but not for avenins, as reported in the literature (Fig. 1A). In contrast, the GLD7 antibody (the recognised epitope is not specified by the manufacturer) detected proteins in all preparations, indicating the presence of conserved prolamin epitopes shared between wheat and oats (Fig. 1B). This suggests partial antigenic overlap despite compositional differences.

**Figure 1:**
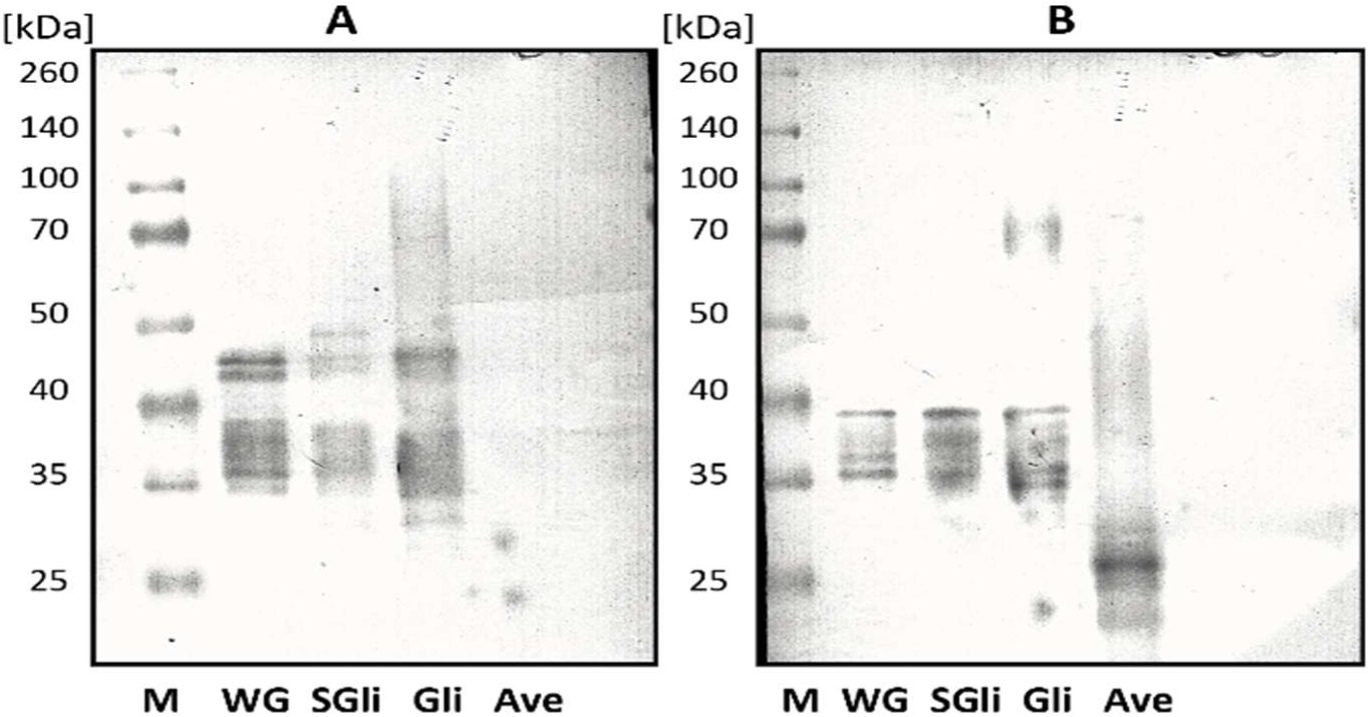
Identification of isolated proteins by western blot. (A) Vital wheat gluten (VG), commercial gliadin (SGli), isolated gliadins (Gli) and isolated avenins (Ave) detected with the G12 antibody. (B) The same preparations were detected with the GLD7 antibody. M, molecular-mass marker.

### 3.2. Effects of prolamin-enriched diets on general growth parameters

To evaluate the impact of dietary prolamin supplementation on systemic development, body weight and morphometric parameters were assessed following the five-week intervention. Final body weights, organ masses, and colon lengths showed no significant deviations between groups, suggesting that the experimental diets were well-tolerated and did not induce systemic toxicity or maladaptive nutritional status.

However, specific morphometric changes were observed in the treatment groups. Mice receiving the avenin-enriched diet exhibited a modest but statistically significant reduction in body length (Fig. 2A; AVE 9.07 cm vs. 9.48 cm in CTRL; p < 0.05), accompanied by a trend toward shorter small intestinal length that approached statistical significance (Fig. 2A; AVE 38.27 cm vs. 40.25 cm in CTRL). Concurrently, the gliadin-enriched group demonstrated a significantly lower cumulative weight gain over the course of the experiment (Fig. 2A; GLI 0.58 g vs. 2.06 g in CTRL; p < 0.01).

**Figure 2:**
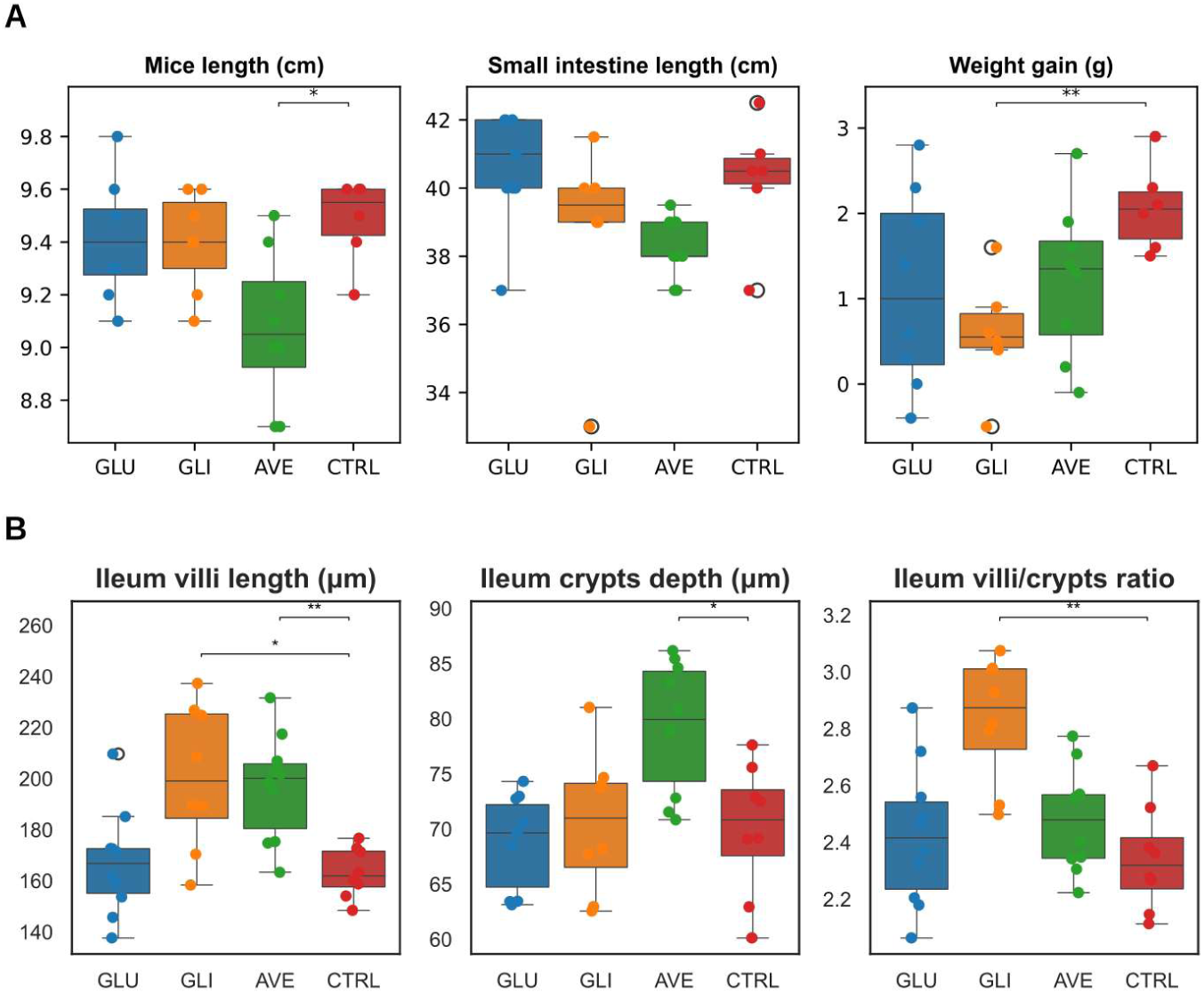
General animal parameters and ileal villus and crypt morphometry after five weeks of dietary intervention. (A) Mice fed the AVE diet exhibited reduced body length, and the GLI group showed lower cumulative body-weight gain. A shorter small-intestinal length was observed in AVE as a non-significant trend. Asterisks indicate significant comparisons shown in the panels. (B) GLI and AVE groups showed longer ileal villi than CTRL; AVE additionally showed increased crypt depth, and GLI showed a higher villus-to-crypt ratio. Asterisks indicate significant post-hoc comparisons displayed in the panels.

### 3.3. Small intestinal morphology and region-specific adaptation

To assess the impact of dietary prolamin intake on mucosal architecture, villus length and crypt depth were quantified across the duodenum, jejunum, and ileum following established methodology. Histological evaluation revealed a normal small bowel with fully maintained epithelial structural integrity, showing no evidence of inflammation, villous atrophy, crypt hyperplasia, or erosive lesions across any segment (Supplementary Data D). Additionally, Peyer’s patches in the ileal segment were clearly visible without significant lymphocyte hyperplasia or enlargement, collectively confirming the absence of overt histopathological damage.

Significant morphometric alterations were localized specifically to the ileal mucosa. Compared to controls, ileal villus length was significantly increased in mice supplemented with either gliadin (Fig. 2B; GLI 200.63 µm vs. CTRL 163.22 µm; p < 0.05) or avenin (Fig. 2B; AVE 196.80 µm vs. CTRL 163.22 µm; p < 0.01). Furthermore, the avenin-fed group exhibited a concomitant increase in crypt depth (Fig. 2B; AVE 79.35 µm vs. CTRL 69.99 µm; p < 0.05). Consequently, the villus-to-crypt ratio was significantly elevated in the gliadin group (Fig. 2B; GLI 2.8 µm vs. CTRL 2.3 µm; p < 0.01).

### 3.4. Humoral responses to dietary prolamins

To evaluate systemic humoral responses to dietary antigens, plasma IgG and IgA reactivity against gluten, gliadin, avenin, and casein were quantified.

In mice receiving the gluten-supplemented diet, plasma IgG levels did not increase and remained comparable to those of the control group (Fig. 3A and 3B). In contrast, gliadin-fed animals exhibited significantly increased IgG reactivity against both gluten (Fig. 3B; p < 0.05) and gliadin (Fig. 3B; p < 0.01), alongside detectable cross-reactivity with avenin (Fig. 3B; p < 0.05), although the response was heterogeneous and largely driven by two strong responders, with most animals remaining close to baseline levels. The avenin-supplemented diet elicited the broadest IgG response, characterised by significantly elevated reactivity against all tested cereal prolamins (Fig. 3B; p < 0.0001). The absence of detectable IgG reactivity towards casein across all experimental groups confirmed that the humoral response was antigen specific. In contrast to IgG reactivity, plasma IgA levels remained at baseline across all experimental cohorts, with no statistically significant differences observed between dietary groups.

**Figure 3:**
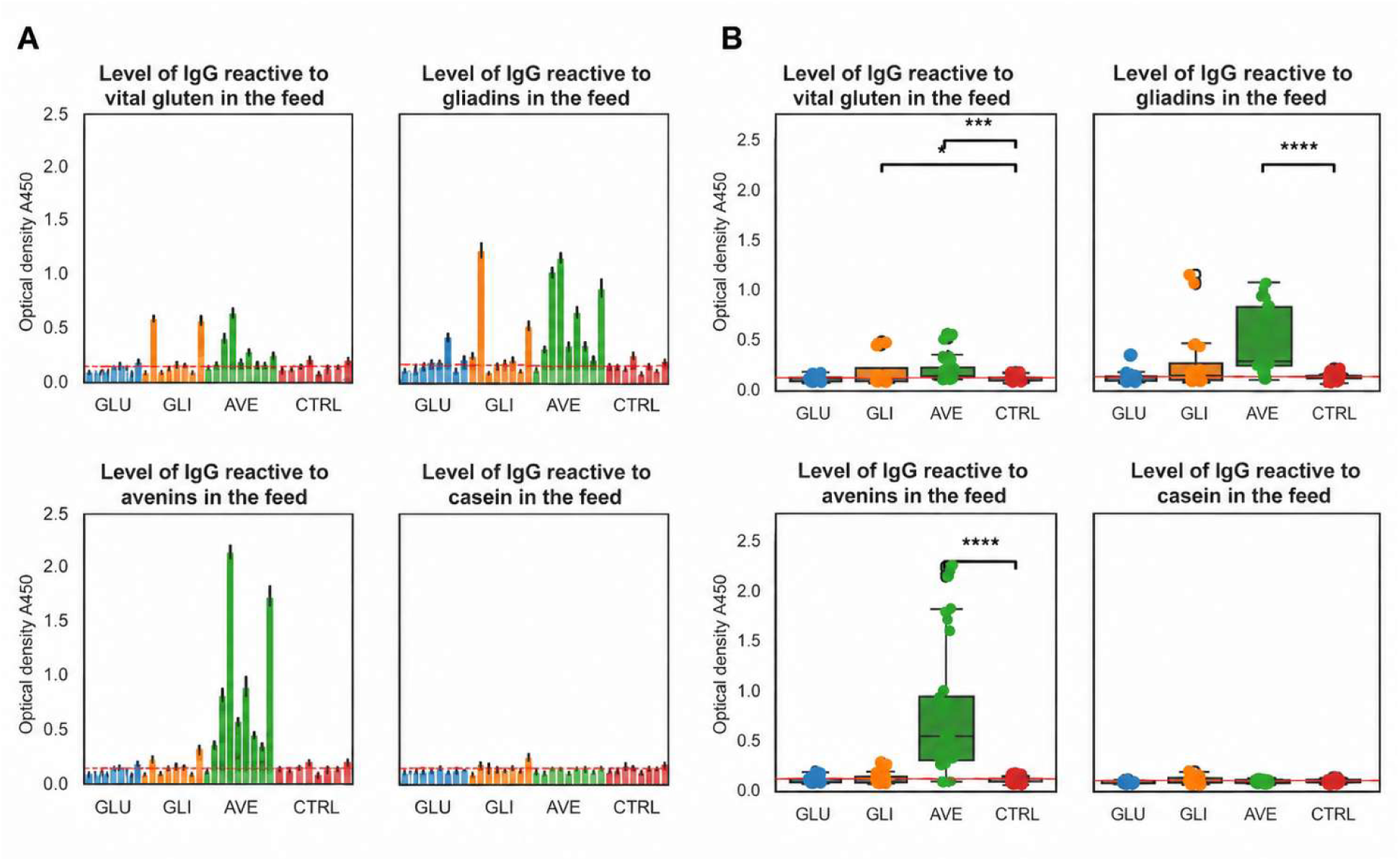
Individual plasma IgG reactivity against gluten, gliadin, avenin and casein. (A) Each bar represents one mouse at plasma dilution 1:250. The orange horizontal reference indicates the mean CTRL response for the corresponding antigen. AVE animals displayed the broadest and most heterogeneous cereal-prolamin reactivity. (B) Group-level plasma IgG reactivity against gluten, gliadin, avenin and casein. GLI animals showed increased IgG reactivity to gluten, gliadin and avenin, whereas AVE animals showed the strongest responses across all three cereal prolamins. No corresponding elevation was observed against casein. Plasma dilution 1:250.

### 3.5. Selective reduction in IL-10 expression correlates with humoral responsiveness

To determine whether dietary prolamins elicited overt mucosal inflammation, transcriptional profiling of key cytokines associated with innate and adaptive immunity was performed. Expression levels of IL-1ß, IL-2, IL-13, IL-15, IL-17a, IL-21, TNF-α, and IFN-γ remained indistinguishable among all experimental groups, confirming the absence of generalized inflammatory activation. Similarly, transcripts essential for epithelial barrier integrity, including tight junction proteins (Tjp1, Tjp2, Cldn2, Cldn15, and Ocln), showed no significant changes, suggesting that barrier function was preserved throughout the intervention.

Because the humoral response showed a continuous and heterogeneous distribution, a post hoc exploratory IgG-phenotype analysis was performed. Animals with antigen-specific IgG responses exceeding the mean response of the control group were classified as responders (Positive), whereas animals with responses at or below this threshold were classified as non-responders (Negative). The phenotype-based analysis revealed a selective reduction in IL-10 expression in IgG-positive animals (Fig. 4; p < 0.001), suggesting that the observed change was associated more closely with humoral responsiveness than with treatment-group assignment alone. This biological relationship was not apparent when animals were analysed solely according to their dietary groups.

**Figure 4:**
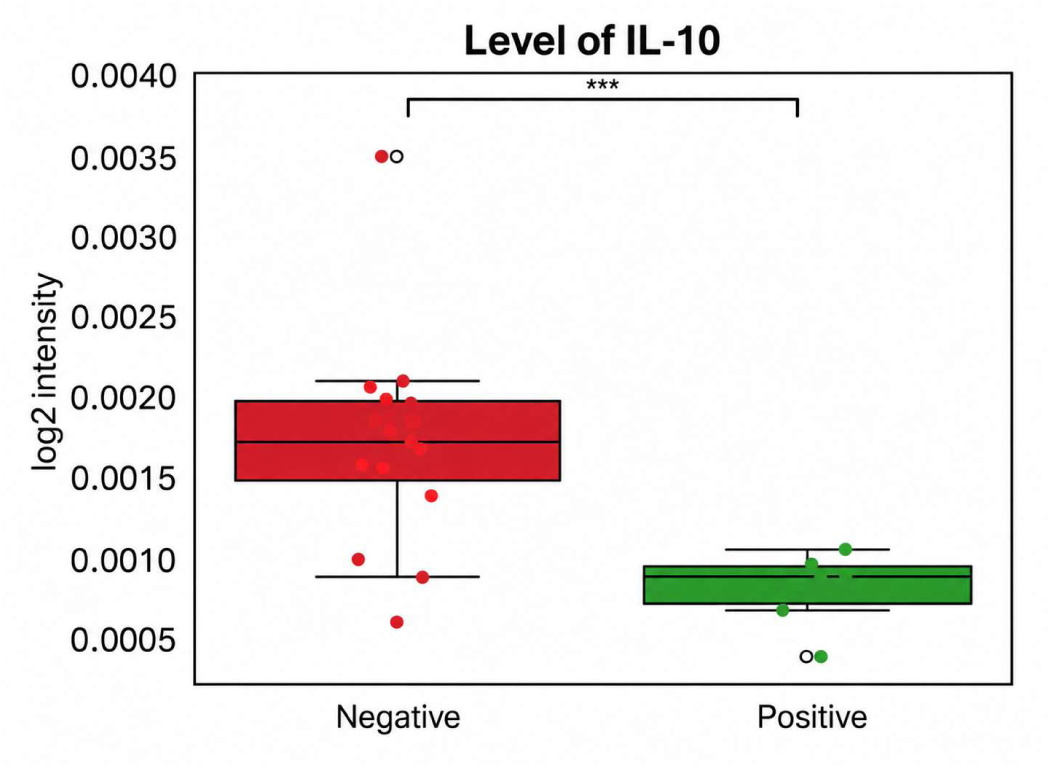
Mucosal IL-10 expression according to antigen-specific IgG responder phenotype. Animals classified as IgG-positive responders showed lower IL-10 expression than IgG-negative animals. This post hoc phenotype-based analysis was used for hypothesis generation rather than as a diet-specific confirmatory comparison.

### 3.6 Descriptive proteomic profiling revealed diet-associated functional patterns

Proteins included in the descriptive proteomic overview were assigned to seven composite GO-derived categories: Extracellular Matrix and Adhesion; Luminal Proteolysis; Immunity, Stress and Cell Survival; Energy and Lipid Metabolism; Membrane and Apical Transport; Nucleus and Gene Regulation; and Cytosol and Cytoskeleton (Fig. 5). GLU showed changes distributed across functional categories, without any single category dominating the overall profile. GLI was characterised by increases in Luminal Proteolysis (18%) and Extracellular Matrix and Adhesion (16%), accompanied by smaller changes in Energy and Lipid Metabolism and Membrane and Apical Transport (13% each). In contrast, AVE exhibited predominantly downward changes in Nucleus and Gene Regulation (19%), Extracellular Matrix and Adhesion (18%), and Cytosol and Cytoskeleton (17%). These protein categories are commonly altered during intestinal maturation; proteomic studies have associated epithelial differentiation with changes in brush-border organisation, lipid uptake and metabolism, and cytoskeletal proteins (Chang et al., 2008). Similarly, dietary intervention studies in healthy mice have further shown that protein source influences ileal transcriptional profiles and nutrient-responsive signalling (Kar et al., 2017).

**Figure 5:**
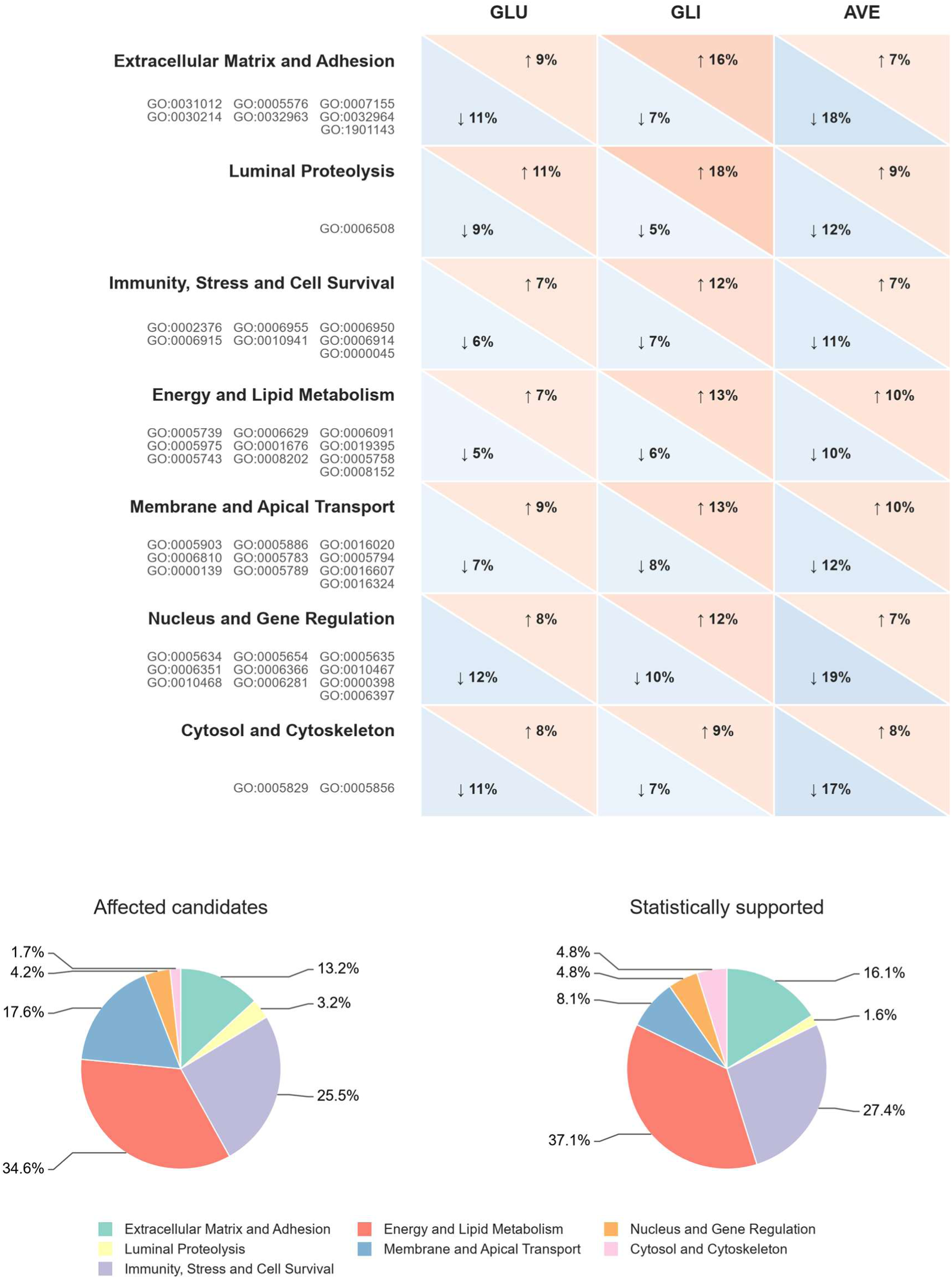
Global proteomic overview using seven composite GO-derived functional categories. The upper matrix shows the descriptive proportions of upward and downward changes in GLU, GLI and AVE relative to CTRL. Pie charts compare the distribution of affected candidates and the statistically supported subset across the same categories. Percentages represent the relative contribution of each composite category.

The functional distribution of the broader affected-candidate dataset and the statistically supported subset was largely preserved, with Energy and Lipid Metabolism, Immunity, Stress and Cell Survival, and Extracellular Matrix and Adhesion remaining the most represented categories. Membrane and Apical Transport showed lower representation in the statistically supported subset (17.6% vs. 8.1%; Fig. 5).

Related functional areas have also been identified in human intestinal transcriptomic studies of diseases associated with gluten intolerance. Sowińska et al. (2020) identified GO terms associated with plasma-membrane and extracellular compartments, whereas Bragde et al. (2018) reported enriched GO categories encompassing membrane and extracellular components, metabolism, transport and immune processes. These findings provide a broader biological context for the functional categories observed in our study, without implying equivalent changes in individual proteins (Bragde et al., 2018; Sowińska et al., 2020). The statistically supported subset comprised 62 proteins, classified by directional pattern as common (26), GLU/GLI-shared (12), GLU-predominant (4), GLI-predominant (13), and AVE-predominant (7). Individual abundance ratios and statistical comparisons are provided in Supplementary Table S1.

### 3.7 Shared structural and metabolic responses

Three fibrillar collagens, COL3A1, COL1A1 and COL1A2 (full names of all proteins are provided in the Supplementary Data), showed increased abundance across all three dietary groups, with greater increases in COL1A1 and COL1A2 in GLU and GLI than in AVE (COL3A1: ANOVA p = 0.013; adjusted p = 0.010, 0.026 and 0.040; COL1A1: ANOVA p = 0.033; adjusted p = 0.037, 0.037 and 0.293; COL1A2: ANOVA p = 0.020; adjusted p = 0.014, 0.014 and 0.186 for GLU, GLI and AVE, respectively; Fig. 6). As major components of the intestinal extracellular matrix, collagens provide structural support and contribute to epithelial growth, differentiation and migration (Pompili et al., 2021; Schuppan & Riecken, 1990). Their increased abundance accompanied longer ileal villi in GLI and AVE, together with deeper crypts in AVE and a higher villus-to-crypt ratio in GLI, suggesting a possible contribution of ECM remodelling to the observed changes in mucosal architecture. Alongside these structural changes, two mitochondrial proteins, GLS and TMEM186, were increased across all three dietary groups (GLS: ANOVA p = 0.0067; adjusted p = 0.046, 0.0028 and 0.038; TMEM186: ANOVA p = 0.0051; adjusted p = 0.0031, 0.0051 and 0.049 for GLU, GLI and AVE, respectively; Fig. 6). GLS converts glutamine to glutamate, an important metabolic substrate for enterocytes, and its increased abundance may be related to the glutamine-rich composition of cereal prolamins (Cruzat et al., 2018; Wieser, 2007). TMEM186 contributes to mitochondrial respiratory complex I assembly (Formosa et al., 2020), and its parallel increase may reflect adjustments in mitochondrial energy metabolism.

**Figure 6:**
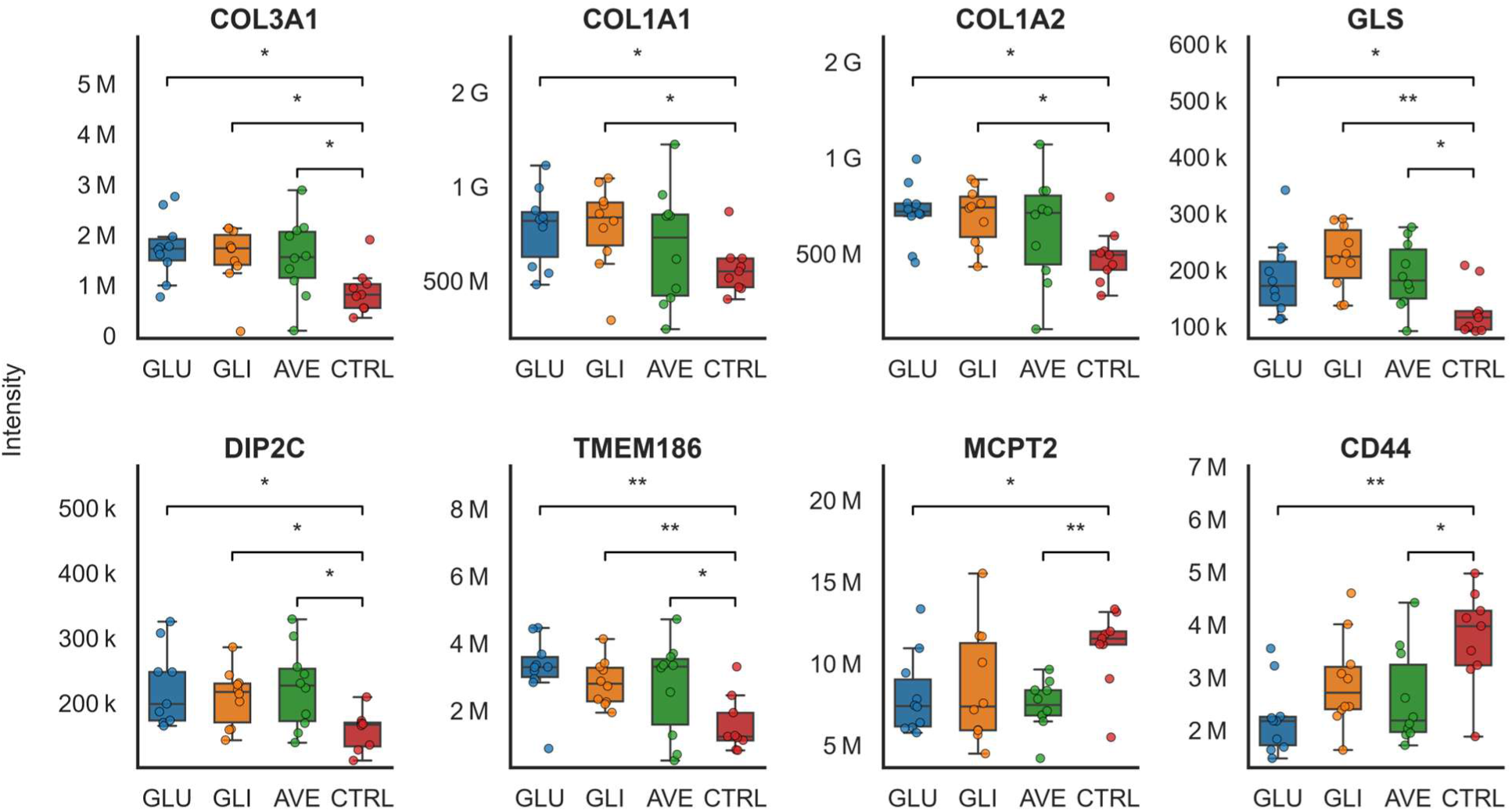
Selected shared protein responses. Upper panels show COL3A1, COL1A1, COL1A2 and GLS; lower panels show DIP2C, TMEM186, MCPT2 and CD44. Box plots and individual data points show abundance in GLU, GLI, AVE and CTRL; brackets indicate the comparisons displayed in each panel.

The shared response also included reduced abundance of MCPT2 and CD44, with the lowest CD44 levels observed in GLU and AVE (MCPT2: ANOVA p = 0.016; adjusted p = 0.038, 0.100 and 0.0069; CD44: ANOVA p = 0.0078; adjusted p = 0.0033, 0.060 and 0.031 for GLU, GLI and AVE, respectively; Fig. 6). Reduced abundance of MCPT2, a mucosal mast-cell protease, may reflect changes in mast-cell abundance or protease expression (Serafin et al., 1990). CD44 is a cell surface glycoprotein found on lymphoid and epithelial cells that contributes to cell–matrix interactions and has been implicated in epithelial organisation and regeneration (Kennel et al., 1993; Walter et al., 2022).

Further changes in intracellular homeostasis involved increased abundance of DIP2C, GET1 and IYD across all three dietary groups, with the highest levels observed in AVE (DIP2C: ANOVA p = 0.010; adjusted p = 0.032, 0.032 and 0.032; GET1: ANOVA p = 0.040; adjusted p = 0.051, 0.051 and 0.035; IYD: ANOVA p = 0.0049; adjusted p = 0.054, 0.0073 and 0.0074 for GLU, GLI and AVE, respectively). These proteins have been linked to lipid regulation, membrane-protein biogenesis and metabolite recycling, respectively (Carvalho et al., 2019; Chai et al., 2025; Thomas et al., 2009), although their roles in the observed intestinal response remain unclear. EXOSC1/CSL4 showed the opposite direction, decreasing across all three diets, particularly in GLU (ANOVA p < 0.001; adjusted p < 0.001, 0.019 and 0.054 for GLU, GLI and AVE, respectively). INTS14 increased most prominently in GLU, with smaller increases in GLI and AVE (ANOVA p < 0.001; GLU adjusted p = 0.0053), indicating a stronger response in GLU rather than an exclusively GLU-associated change.

### 3.8 Wheat-associated digestion, metabolic and epithelial responses

Both GLU and GLI showed increased abundance of pancreatic digestive enzymes, including carboxypeptidases A1 and B1, elastase 3A, chymotrypsin C and trypsin, with greater increases in GLI. Their higher abundance may reflect altered luminal persistence or increased digestive enzyme input into the intestinal lumen. Previous studies reporting increased pancreatic chymotrypsinogen synthesis following gluten feeding and resistance of selected gliadin peptides to pancreatic and brush-border proteolysis provide a possible explanation for these changes (Hausch et al., 2002; Johnson et al., 1977; Shan et al., 2002).

The wheat-associated response also involved proteins related to lipid metabolism and biotransformation (Fig. 7). ACOT1, which regulates fatty-acid availability by hydrolysing fatty acyl-CoA, increased in both GLU and GLI (ANOVA p < 0.001; adjusted p < 0.001 and 0.039, respectively). A similar pattern was observed for CYP2B10 and E9Q593/CYP2B23, while CYP2J6 and CYP4F3 followed the same descriptive trend, with the highest abundance in GLU, intermediate levels in GLI and the lowest levels in AVE. CYP3A25 differed from this pattern, showing increases in GLU and AVE and a smaller increase in GLI (ANOVA p = 0.021; adjusted p = 0.017, 0.228 and 0.043 for GLU, GLI and AVE, respectively). These findings suggest a predominantly wheat-associated response involving fatty-acid handling and biotransformation, alongside additional changes shared by GLU and AVE.

**Figure 7:**
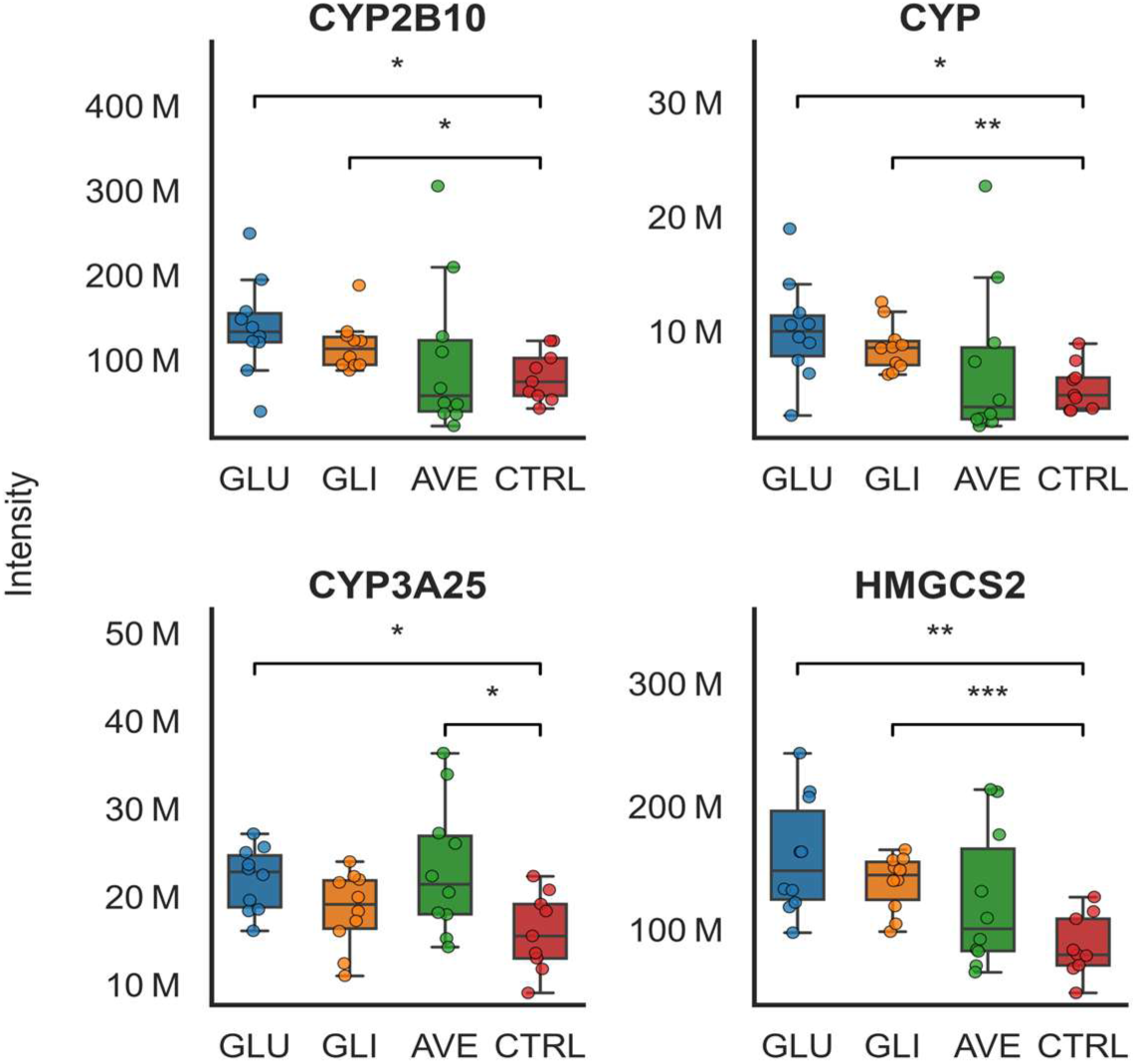
Selected proteins associated with biotransformation and nutrient metabolism. The supplied panels show CYP2B10, an additional cytochrome P450 protein (labelled CYP), CYP3A25 and HMGCS2 in GLU, GLI, AVE and CTRL. Individual mouse intensities, group box plots and indicated statistical comparisons are displayed.

HMGCS2, a mitochondrial enzyme involved in ketogenesis and epithelial differentiation through β-hydroxybutyrate signalling, also increased predominantly in GLU and GLI (Wang et al., 2017; ANOVA p < 0.001; adjusted p = 0.0018, < 0.001 and 0.086 for GLU, GLI and AVE, respectively; Fig. 7). GLU showed a further increase in ADIPOR2, a receptor involved in lipid and energy metabolism, whereas its abundance remained close to CTRL in GLI and AVE (ANOVA p < 0.001; GLU adjusted p < 0.001). Additional GLU-associated changes included increased ABHD17B, involved in the reversible lipid modification of membrane proteins, and altered mitochondrial proteins, with increased FASTKD5 and reduced SDHAF2 (ABHD17B: ANOVA p = 0.0035, adjusted p = 0.042; FASTKD5: ANOVA p = 0.0058, adjusted p = 0.017; SDHAF2: ANOVA p = 0.041, adjusted p = 0.018). Together, these proteins extend the metabolic response from lipid handling and biotransformation to mitochondrial regulation, with additional changes particularly evident in GLU.

GLU and GLI also shared changes in proteins involved in cell-death regulation (Fig. 8). FAS abundance increased modestly in both groups, accompanied by higher abundance of its inhibitory regulator BFAR and reduced BID, which links caspase-8-dependent death-receptor signalling to mitochondrial apoptosis (Li et al., 1998; Zhang et al., 2000; FAS: ANOVA p = 0.015, GLI adjusted p = 0.041; BFAR: ANOVA p = 0.016, GLU adjusted p = 0.0087; BID: ANOVA p = 0.041, GLI adjusted p = 0.045). The accompanying increase in BFAR and reduction in BID may represent compensatory regulation associated with the modest rise in FAS abundance.

**Figure 8:**
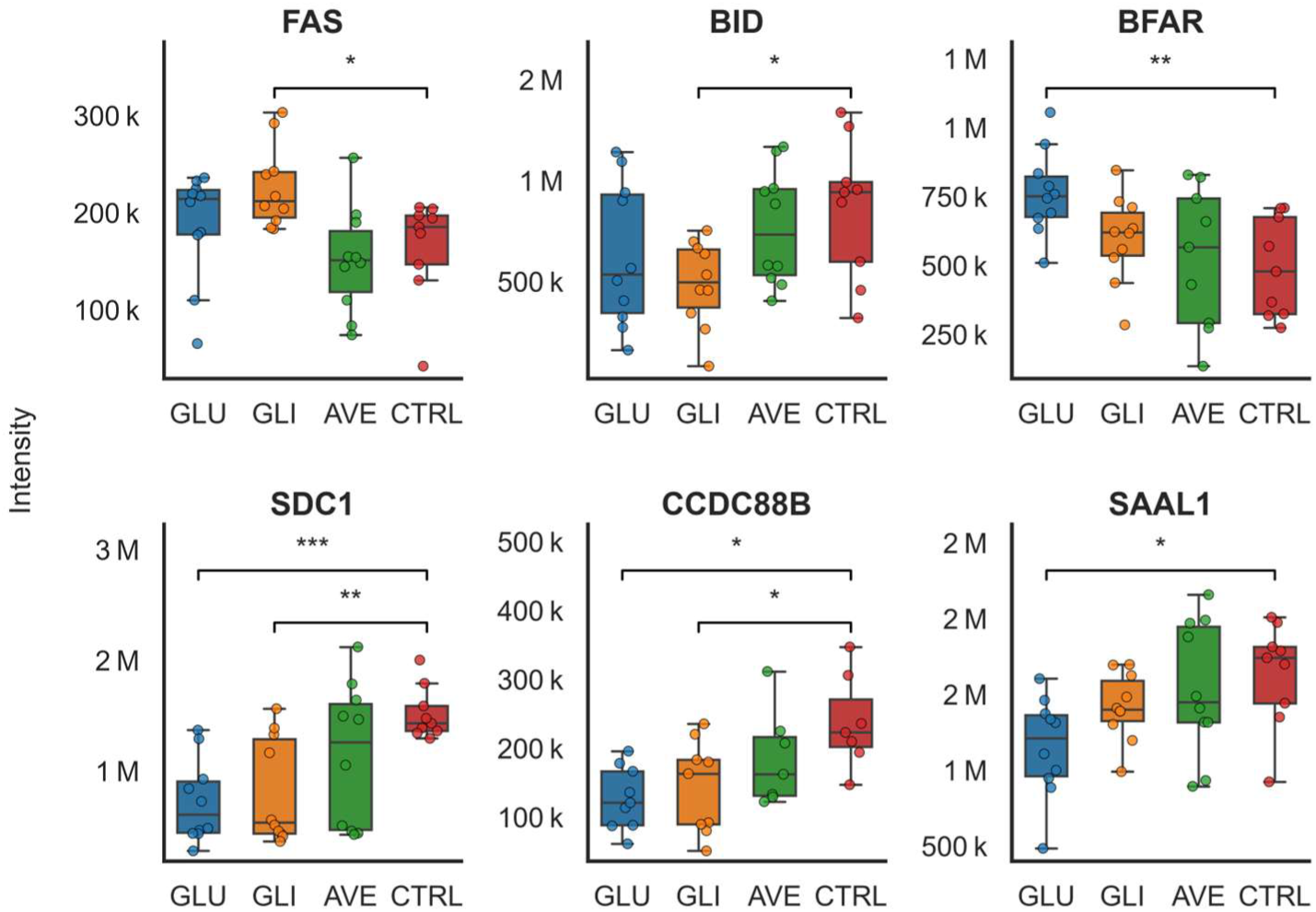
Wheat-associated changes in protein abundance. Upper panels show proteins involved in cell-death regulatory proteins. The lower panels show proteins associated with the epithelial surface and tissue organization. SDC1 and CCDC88B were reduced in both wheat-derived groups, while SAAL1 showed lower abundance particularly in GLU. The individual mouse intensities, group box plots and indicated statistical contrasts are displayed.

Changes in epithelial-surface organisation were reflected by a marked reduction in SDC1 in both wheat-derived groups (ANOVA p < 0.001; adjusted p < 0.001 and 0.0023 for GLU and GLI, respectively; Fig. 8). Syndecan-1 and epithelial heparan sulphate contribute to intestinal barrier organization (Bode et al., 2008). The reduction in SDC1 also accompanied the increased abundance of fibrillar collagens and lower CD44 levels described above. Syndecan-1 can bind type I and III collagens, whereas CD44 participates in hyaluronan-dependent cell–matrix interactions and intestinal epithelial regeneration (San Antonio et al., 1994; Walter et al., 2022; Woods, 2001). Together, these changes suggest possible alterations in epithelial–matrix organisation, although their functional consequences remain to be established. The reduction in SDC1 occurred without detectable differences in the analysed tight-junction transcripts. CCDC88B and SAAL1 also showed reduced abundance in GLU and GLI (CCDC88B: ANOVA p = 0.019; adjusted p = 0.013 and 0.033 for GLU and GLI, respectively).

Further changes involved proteins associated with epithelial sensing and intracellular signalling. TRPV4 increased in both wheat-derived groups, particularly in GLU (ANOVA p = 0.0059; adjusted p = 0.0061 and 0.150 for GLU and GLI, respectively). TRAF6 showed a greater increase in GLI, while ZDHHC21 decreased in both groups, again more prominently in GLI (TRAF6: ANOVA p = 0.0116; adjusted p = 0.018 and 0.725 for GLI and GLU, respectively; ZDHHC21: ANOVA p = 0.0105; adjusted p = 0.0076 and 0.415 for GLI and GLU, respectively). GLU additionally showed increased STAT5A and PLCG1, proteins involved in cytokine-responsive and receptor-dependent intracellular signalling (STAT5A: ANOVA p = 0.023, adjusted p = 0.010; PLCG1: ANOVA p = 0.033, adjusted p = 0.033). These changes indicate that the wheat-associated response extended beyond metabolism to epithelial sensing and intracellular regulation, with differences in the proteins affected by GLU and GLI.

### 3.9 Gliadin-associated sphingolipid and junctional responses

Sphingolipid metabolism formed a further component of the GLI response, with increased abundance of SGPP1 and ASAH2, enzymes involved in sphingosine-1-phosphate and ceramide turnover, respectively (Kono et al., 2006; Le Stunff et al., 2002; SGPP1: ANOVA p = 0.024, adjusted p = 0.023; ASAH2: ANOVA p = 0.021, adjusted p = 0.022). Changes in cytoskeletal and junctional regulation accompanied this metabolic profile. MYL4 increased, whereas ARHGEF18/p114RhoGEF, a regulator of junction-associated RhoA activation and epithelial junction formation, decreased (Terry et al., 2011; MYL4: ANOVA p = 0.030, adjusted p = 0.049; ARHGEF18: ANOVA p = 0.015, adjusted p = 0.0055).

The remaining GLI-associated changes encompassed cell-cycle control, immune signalling, redox metabolism and endothelial function. CDK1, IRF4, ITPKC, PLEK and NADK were reduced (CDK1: ANOVA p = 0.034, adjusted p = 0.031; IRF4: ANOVA p = 0.019, adjusted p = 0.020; ITPKC: ANOVA p = 0.027, adjusted p = 0.045; PLEK: ANOVA p = 0.049, adjusted p = 0.035; NADK: ANOVA p = 0.0025, adjusted p = 0.0093), while CD300LG increased (ANOVA p = 0.024, adjusted p = 0.013; Supplementary Table S1). Their combined biological significance remains unclear.

### 3.10 Avenin-associated epithelial and regulatory responses

The AVE-associated profile included marked reductions in PTPRH/SAP-1 and IL-18, two proteins involved in epithelial and immune regulation (PTPRH: ANOVA p = 0.015, adjusted p = 0.028; IL-18: ANOVA p = 0.044, adjusted p = 0.034; Fig. 9). PTPRH localises to epithelial brush-border microvilli, and experimental studies have linked its loss to altered epithelial immune signalling (Murata et al., 2015; Sadakata et al., 2009). The effects of IL-18 likewise depend on its cellular source and abundance (Harrison et al., 2015; Nowarski et al., 2015). These proteomic changes accompanied longer ileal villi, deeper crypts and frequent antigen-specific IgG responses, suggesting that changes in mucosal regulation or antigen handling may connect the structural, proteomic and humoral findings.

**Figure 9:**
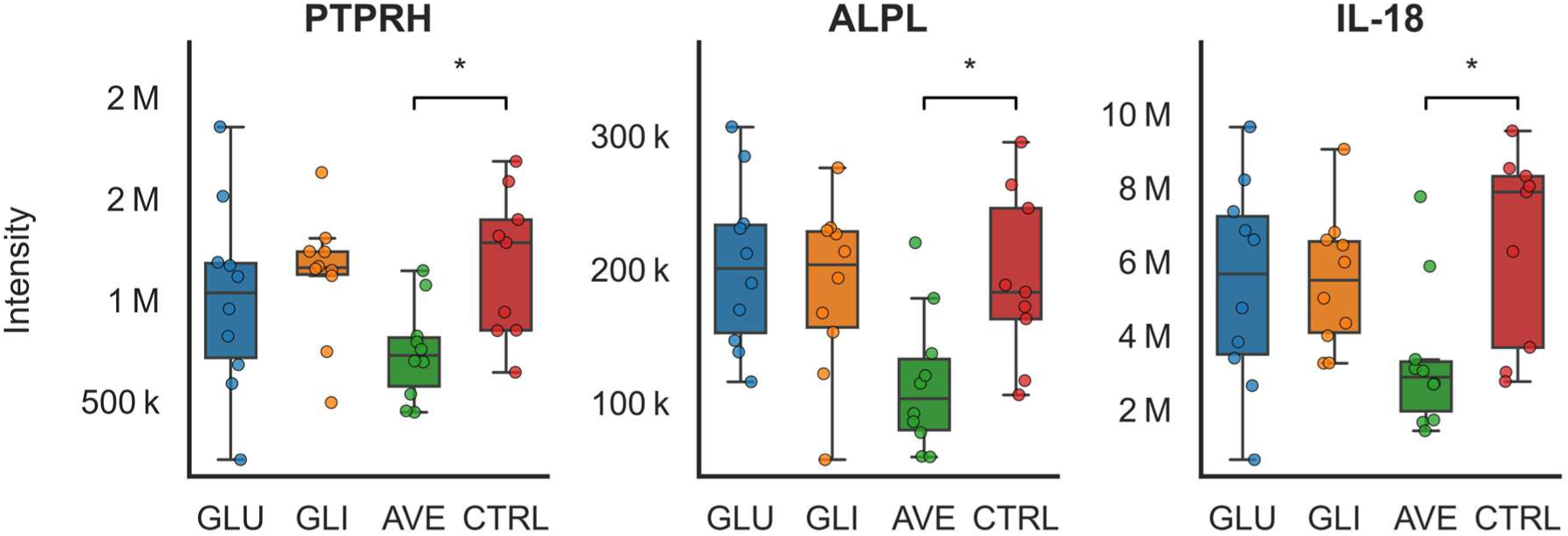
Selected avenin-associated protein responses. PTPRH, ALPL and IL-18 show lower abundance in AVE relative to CTRL; individual mouse intensities, group box plots and indicated statistical comparisons are displayed.

Further proteins affected in AVE include ALPL, HYPK, PRCC and KIF15 (ALPL: ANOVA p = 0.011, adjusted p = 0.033; HYPK: ANOVA p = 0.031, adjusted p = 0.016; PRCC: ANOVA p = 0.013, adjusted p = 0.023; KIF15: ANOVA p = 0.014, adjusted p = 0.016; Fig. 9). PRCC also showed lower abundance in GLI (adjusted p = 0.222), whereas MED24 increased in AVE (ANOVA p = 0.032, adjusted p = 0.037; Supplementary Table S1). The relationship of these changes to the epithelial and humoral findings remains unclear.

### 3.11 Integration and limitations

Together, the findings indicate partly shared intestinal responses to the three dietary preparations, accompanied by modest changes in ileal morphology and antigen-specific IgG reactivity, particularly in GLI and AVE, without overt histopathological injury. Shared properties of cereal prolamins may contribute to the common proteomic response, while differences in the composition and digestion of the preparations, their antigenic properties and host regulation may influence the diet-specific profiles.

Differences in gastrointestinal processing could help explain why the extent of proteomic changes did not parallel the frequency of IgG responses. Persistent wheat-derived peptides may influence luminal digestive processes, whereas the distinct sequence composition of avenins could produce a different pool of peptides and amino acids at the epithelial surface (Gilissen et al., 2016). Differences in the availability and handling of these digestion products may influence nutrient utilisation and antigen exposure, potentially connecting the AVE-associated epithelial and immune-regulatory changes with its more frequent humoral response. However, the mechanisms of digestion, antigen uptake and subsequent immune recognition were not directly examined.

The common ECM changes, wheat-associated SDC1 reduction, GLI-associated ARHGEF18/MYL4 pattern and AVE-associated PTPRH decrease involve proteins associated with epithelial organisation and maintenance. Related functional processes have been identified in human intestinal transcriptomic studies examining the impact of gluten, including extracellular-matrix organisation, adhesion, enterocyte metabolism, nutrient transport and epithelial differentiation (Bragde et al., 2018; Dotsenko et al., 2021, 2024; Loberman-Nachum et al., 2019; Ramírez-Sánchez et al., 2025; Sowińska et al., 2020). These parallels concern shared biological processes rather than equivalent molecular responses or tissue outcomes. Dietary protein composition may also influence intestinal immune-cell populations and antigen-specific repertoires without overt tissue injury (Lockhart et al., 2023). In a related study in BALB/c mice, five-week diets with equivalent inclusion of purified gluten, gliadin or avenin produced no histopathological changes in the ovaries, testes or thyroid gland, although morphometric differences, mainly in follicle, oocyte and seminiferous tubule dimensions, were more pronounced in the gluten-and gliadin-fed groups than in the avenin and gluten-free groups, and were considered to fall within the range of normal physiological adaptation (Makovický et al., 2026).

Although dietary exposure was experimentally controlled, the relationships between the proteomic, morphological and humoral findings remain observational. The overall phenotypic changes were modest, as might be expected in otherwise unsensitive BALB/c mice, and the present findings primarily identify candidate molecular responses rather than establish their functional consequences. In particular, mucosal secretory IgA, cellular immune responses, intestinal permeability and the potential contribution of diet-associated microbiome changes were not assessed. Further studies combining targeted protein validation, cellular localisation and functional measurements will be needed to determine whether the observed changes represent physiological adaptation to dietary prolamins, compensatory responses or processes with longer-term consequences.

## 4. Conclusion

Equivalent dietary inclusion of gluten, purified gliadin and purified avenin produced modest but distinct intestinal responses in BALB/c mice without overt histopathological injury. All three diets shared changes in extracellular-matrix proteins, glutamine-related metabolism, mitochondrial proteins and mucosal regulation. Increased fibrillar collagens accompanied longer ileal villi in GLI and AVE, while reductions in CD44 across the dietary groups and SDC1 in the wheat-derived groups suggest possible changes in epithelial–matrix interactions and mucosal organisation. The parallel increases in GLS and TMEM186 further indicate a shared response involving proteins associated with glutamine utilisation and mitochondrial energy metabolism.

Beyond these common changes, GLU and GLI showed related responses in digestive enzymes, lipid metabolism and biotransformation, with additional mitochondrial and metabolic changes particularly evident in GLU. GLI also exhibited changes in sphingolipid metabolism and junctional regulation, whereas AVE showed a distinct epithelial and immune-regulatory profile, including reduced PTPRH and IL-18. Despite equivalent dietary inclusion of purified gliadins and avenins, AVE elicited more frequent cereal-prolamin-specific IgG responses, while IgG-positive animals showed lower intestinal IL-10 expression. These findings suggest that the intestinal response to cereal prolamins involves both shared biological processes and differences associated with the individual protein preparations. Given the modest overall phenotypic changes, the observed relationships remain exploratory. Further functional studies are needed to determine how differences in protein digestion, epithelial–matrix organisation and mucosal antigen handling contribute to the distinct responses observed here.

## CRediT authorship contribution statement

Michaela Šťastná: Writing – original draft, Methodology, Investigation, Formal analysis, Data curation. Adam Norek: Methodology, Investigation, Data curation, Formal analysis, Writing – review & editing. Peter Makovický: Investigation. Edita Jeklová: Investigation. Petra Straková: Investigation. Matěj Hrunka: Writing – review & editing. Petr Jabandžiev: Supervision, Project administration, Funding acquisition Writing – review & editing. Lubomír Janda: Conceptualization, Supervision, Project administration, Funding acquisition, Writing – review & editing.

## Supporting information

Supplementary Data A_Total Protein and Fat Content.pdf

Supplementary Data B_ MS Analysis_250526_6597.xlsx

Supplementary Data C_ RT-qPCR gene sequence.xlsx

## Acknowledgments

CIISB, Instruct-CZ Centre of Instruct-ERIC EU consortium, funded by MEYS CR infrastructure project LM2023042 and European Regional Development Fund Project “Innovation of Czech Infrastructure for Integrative Structural Biology” (No. CZ.02.01.01/00/23_015/0008175), is gratefully acknowledged for the financial support of the measurements at the CEITEC Proteomics Core Facility. Computational resources were provided by the e-INFRA CZ project (ID: 90254), supported by MEYS CR.

## Data availability

Data will be made available on request.

## Declaration of generative AI and AI-assisted technologies in the manuscript preparation process

During the preparation of this work, the authors used artificial intelligence tools solely for language editing and translation purposes, including improving grammar, clarity, and readability of the text. All literature and sources cited in this manuscript were identified, verified, and referenced by the authors themselves. All AI-assisted content was subsequently reviewed and edited by the authors, who take full responsibility for the content, accuracy, and integrity of the final published work.

