## Supplementary Data A_Total Protein and Fat Content.pdf for "Dietary Exposure to Purified Gliadins and Avenins: Effects on Intestinal and Immune Responses in BALB/c Mice"

Food Research Institute Prague
Department of Research
Radiová 7, 102 00 Praha 10

### CERTIFICATE

Certificate No.: 3/24 Total page count.: 1 Page No.: 1

Name and address of customer: Mgr. Michaela Šťastná
Výzkumný ústav veterinárního lékařství, v. v. i.
Hudcova 296/70
621 00 Brno

Sample description and identification:

| | Sample |
|---|---|
| 1 | Wheat gluten |
| 2 | Wheat gliadin |
| 3 | Oat avenin |

Date of sample receipt: March 26, 2024

Method description:

Proteins Kjeldahl method

Fat gravimetrically after acid hydrolysis and chloroform extraction

Result of testing:

| Sample | Nitrogen | Proteins (g/100g) | | Fats |
|---|---|---|---|---|
| | (g/100g) | N x 6,25 | N x 5,7 | (g/100g) |
| gluten | 11,99 | 74,9 | 68,3 | 4,56 |
| gliadin | 15,23 | 95,2 | 86,8 | 0,32 |
| avenin | 14,08 | 88,0 | 80,2 | 0,49 |

Date: April 12, 2024

Authorised person: Ing. Ivana Laknerová
Dept of Research
