## Supplementary Data D.pdf for "Dietary Exposure to Purified Gliadins and Avenins: Effects on Intestinal and Immune Responses in BALB/c Mice"

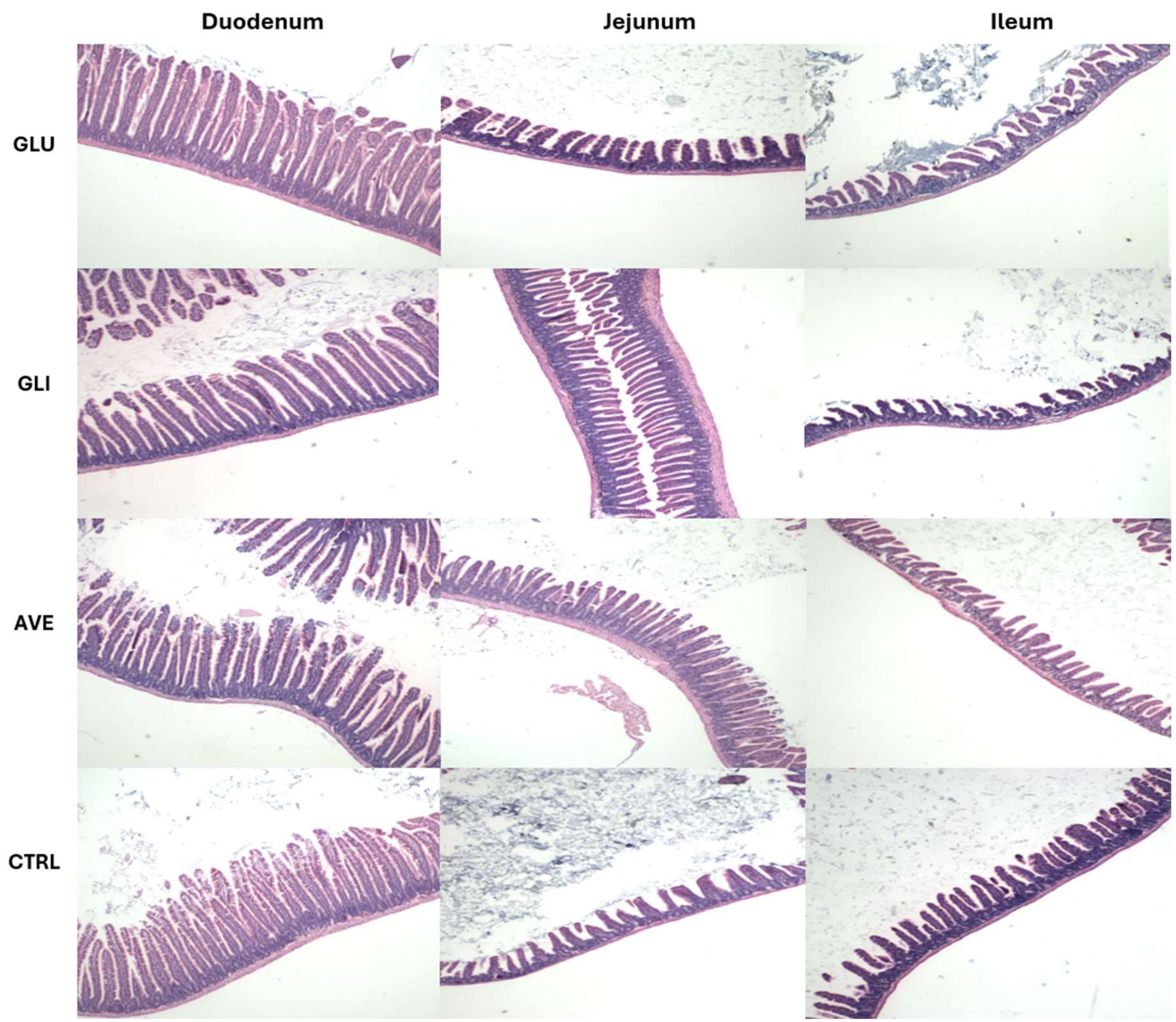


***Figure 1: Histological examination of the small intestine.*** *Representative haematoxylin-eosin-stained duodenal, jejunal and ileal sections from GLU, GLI, AVE and CTRL groups. The mucosa remained structurally intact without villous atrophy, erosive lesions or overt inflammatory pathology.*
